# Reversal of Obesity by Enhancing Slow-wave Sleep via a Prokineticin Receptor Neural Circuit

**DOI:** 10.1101/2024.04.30.591948

**Authors:** Yong Han, Guobin Xia, Lauren Harris, Panpan Liu, Dongyin Guan, Qi Wu

## Abstract

Obese subjects often exhibit hypersomnia accompanied by severe sleep fragmentation, while emerging evidence suggests that poor sleep quality promotes overeating and exacerbates diet-induced obesity (DIO). However, the neural circuit and signaling mechanism underlying the reciprocal control of appetite and sleep is yet not elucidated. Here, we report a neural circuit where prokineticin receptor 2 (PROKR2)-expressing neurons within the parabrachial nucleus (PBN) of the brainstem received direct projections from neuropeptide Y receptor Y2 (NPY2R)-expressing neurons within the lateral preoptic area (LPO) of the hypothalamus. The RNA-Seq results revealed *Prokr2* in the PBN is the most regulated GPCR signaling gene that is responsible for comorbidity of obesity and sleep dysfunction. Furthermore, those NPY2R^LPO^ neurons are minimally active during NREM sleep and maximally active during wakefulness and REM sleep. Activation of the NPY2R^LPO^→PBN circuit or the postsynaptic PROKR2^PBN^ neurons suppressed feeding of a high-fat diet and abrogated morbid sleep patterns in DIO mice. Further studies showed that genetic ablation of the PROKR2 signaling within PROKR2^PBN^ neurons alleviated the hyperphagia and weight gain, and restored sleep dysfunction in DIO mice. We further discovered pterostilbene, a plant-derived stilbenoid, is a powerful anti-obesity and sleep-improving agent, robustly suppressed hyperphagia and promoted reconstruction of a healthier sleep architecture, thereby leading to significant weight loss. Collectively, our results unveil a neural mechanism for the reciprocal control of appetite and sleep, through which pterostilbene, along with a class of similarly structured compounds, may be developed as effective therapeutics for tackling obesity and sleep disorders.

## INTRODUCTION

A growing body of evidence suggests the co-morbidities of obesity and sleep disorders which has led to significant clinical attention recently (*1, 2*). Many of those clinical discoveries were drawn from observations of obstructive sleep apnea, Prader-Willi syndrome, insomnia, as well as narcolepsy where obesity and sleep disorders are taken as complications of each other (*3–10*). So far, there is a lack of effective treatment towards this increasingly common symptom, because the underlying neurological mechanisms are largely elusive. There is accumulating evidence showing that sleep deprivation is associated with obesity (*11, 12*). It is well known that during sleep the secretion of various hormones varies, contributing to the metabolism and energy balance of our body (*13*), which may be disturbed by sleep deprivation. Sleep deprivation decreased leptin levels and increased ghrelin level may explain the change of body weight (*14*).

The parabrachial nucleus (PBN) is traditionally known as an important hub relaying sensory information to the cerebral cortex, basal forebrain, hypothalamus, thalamus, amygdala complex, and descending projections to medullary regions (*15–18*). Recent findings suggest that the PBN in the brainstem plays a major role in regulating multiple physiological activities including feeding and sleep (*19–22*). The PBN neurons expressing GABA_A_ receptors strongly regulate anorexic or appetitive symptoms (*21, 23*). A subgroup of PBN neurons expressing calcitonin gene-related peptide (CGRP) mediates noxious feeding in normal and cachexic models (*24–26*). Meanwhile, the PBN also plays an indispensable role in inducing and maintaining electroencephalogram (EEG) arousal and behavioral wakefulness (*27, 28*). The sleep process is viewed as a process of ‘inhibition of wakefulness’, and thus GABAergic neurons are essential to turning off the signals for arousal or wakefulness (*29–31*).

Notably, wake-promoting neurons such as orexin neurons in the lateral hypothalamus (*32*) and unidentified PBN neurons (*33*), were proved to have GABAergic upstream regulators that functionally contribute to sleep. Clinically, drugs dealing with GABA receptors are most commonly used as hypnotics, and one of the most common adverse reactions of these hypnotics (e.g. benzodiazepine (*34*)) was improved appetite and weight gain (*35, 36*), which is also an intriguing co-occurrence of sleepiness and obesity. However, the neural circuit basis and physiological functions of these PBN neurons are uncharacterized.

While factors that mediate the direct effect of light on sleep behavior are unknown, several factors have been proposed to transmit circadian information from master circadian pacemakers, such as the mammalian suprachiasmatic nucleus (SCN) to other tissues(*37*). One of these factors, prokineticin 2 (PROK2), has been implicated in regulating circadian in nocturnal rodents. *Prok2* mRNA levels oscillate in a circadian manner in the rodent SCN (*38, 39*). The mice that lack PROK2 or PROKR2 exhibit attenuated circadian rhythms of locomotor activity, thermoregulation, and circulating corticosteroid and glucose levels (*40, 41*). These studies suggest a key role for PROK2 in regulating sleep. However, it is still unknown what neural circuits are involved in the sleep regulation by PROK2 signaling.

In this study, we identified a neural circuit that is composed of a subset of neurons in the lateral preoptic area (LPO) that expressed neuropeptide Y receptor Y2 (NPY2R) and PROKR2-expressing neurons in the PBN. Notably, we discovered that pterostilbene, a plant-derived stilbenoid, showed wakefulness-promoting and weight-reducing effects in obese models. Collectively, our results demonstrate a previously unknown circuit for “top-down” septohypothalamic promotion of feeding and sleep, providing new and important insights into the therapy of feeding and sleep disorders.

## RESULTS

### Obesity is comorbid with dysregulated NREM sleep

To ascertain whether obesity and sleep disorders are pathologically associated, we examined the chronic effects of high-calorie diet (HFD, 45 kcal% fat) in diet-induced obesity (DIO) mice on body weight and sleep parameters for 8 weeks. As expected, due to a significant increase of daily calorie intake of HFD, the body weight progressively increased (Fig. 1, A and B). Intriguingly, after 8 weeks of HFD treatment, the DIO mice exhibited significantly decreased wakefulness and increased non-REM (NREM) sleep particularly in nighttime while rapid eye movement (REM) sleep was slightly affected (Fig. 1, C-G). Meanwhile, chronic treatment of HFD for 8 weeks led to a progressive increase in the episode number of NREM sleep and wakefulness, with the episode duration of NREM and wakefulness reduced significantly (Fig. 1, H-K), denoting an emergence of sleep fragmentation. These phenotypes on a large scale corresponded to J.B. Jenkins et al.’s finding(*42*), in which the feeding and sleep parameters were measured in 2 weeks, 4 weeks and 6 weeks of HFD feeding, and elevated body weight and food intake was observed, along with a decline in daily wakefulness, an incline in daily NREM sleep, and mild evidence of sleep fragmentation. In our studies, we also observed a progressive decrease of EEG power during NREM sleep, and specifically, delta power with a frequency between 0.5 and 4 Hz during NREM sleep was significantly decreased in HFD group compared with LFD group (Fig. 1, L-N). Most strikingly, the typical enhancement of delta power during the transition from wakefulness to NREM in WT mice were suppressed with more frequent transitions from wakefulness to NREM stages in DIO mice (Fig. 1, O and P). This evidence demonstrates that HFD induced obesity can demodulate sleep, especially the phenomenal, disrupted wakefulness to NREM transitions and the correspondent decrease of delta power in NREM, which serves as a cause-and-effect link for us to study further on the circuits that regulate sleep and feeding behavior simultaneously.

**Fig. 1.**
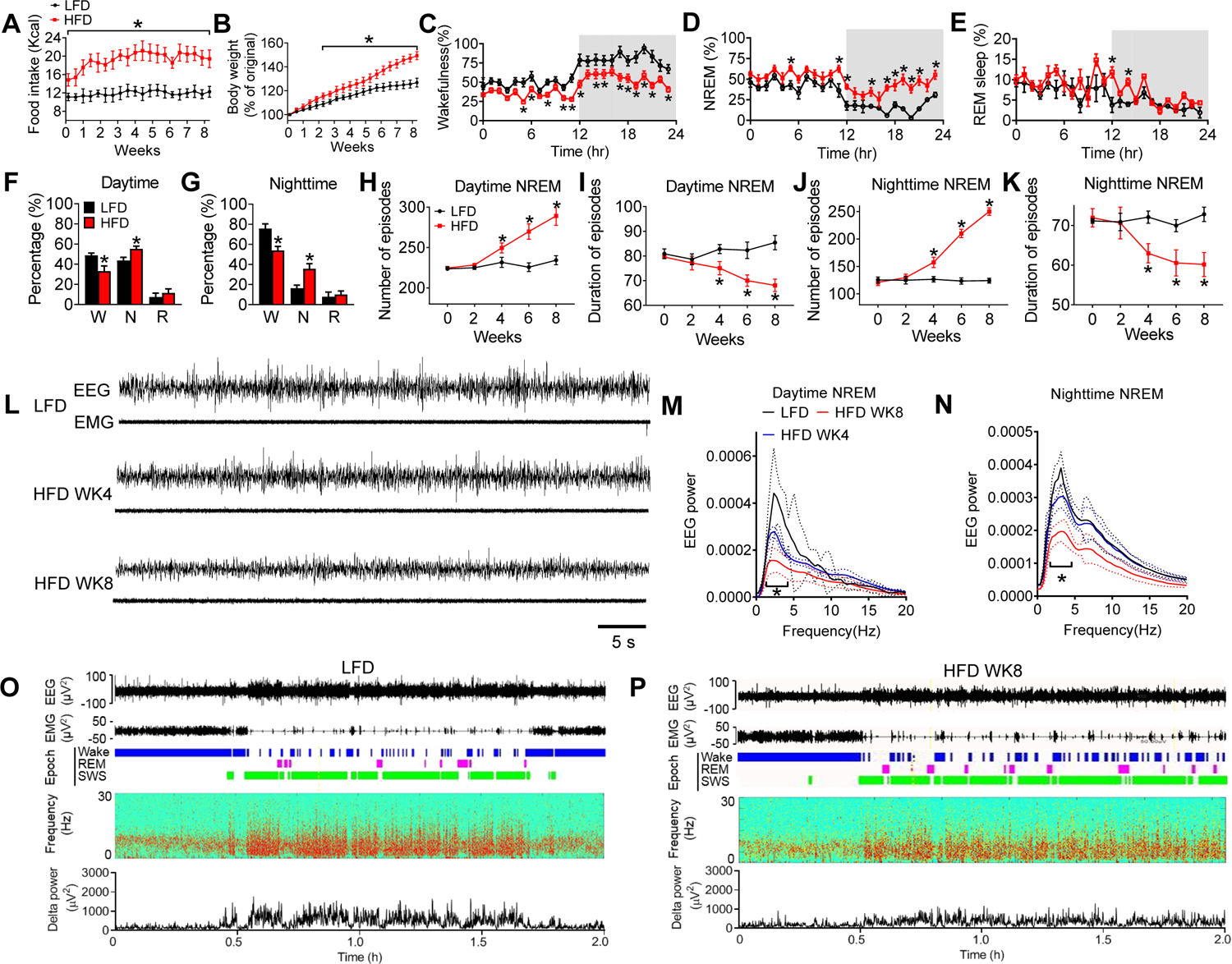
Chronic treatment with high-fat diet induces obesity and sleep disorder. (**A**, **B**) Daily food intake (**A**) and body weight (**B**) monitored for 8 weeks in the mice fed with high-fat diet (HFD). (**C**-**E**)Time spent in wakefulness (**C**), NREM sleep (**D**), and REM sleep (**E**) during the 24-hr period on the day 56 in the mice described in **A**, **B**. (**F**, **G**) Percent of time spent per day in wakefulness, NREM sleep, and REM sleep in the mice described in **C-E**. (**H**-**K**) The number (**H**, **J**) and duration (**I**, **K**) of NREM sleep episodes during daytime and nighttime in the mice described in **A**, **B**. (**l**) Representative EEG and EMG recordings in the mice treated with LFD, HFD for 4 weeks and HFD for 8 weeks. (**M**, **N**) EEG power from NREM sleep during daytime (**M**) and nighttime (**N**) in the mice described in **L**. (**O**, **P**) Representative EEG and EMG traces, brain states (color coded), power spectra, and delta power from EEG on a 2-hr timescale during daytime in the mice treated with LFD (**O**) and HFD for 8 weeks (**P**). Error bars represent mean ± SEM. n = 6-8 per group; *p < 0.05; two-way ANOVA and followed by Bonferroni comparisons test.

### *Prokr2* gene in the PBN is the key to regulating sleep and feeding

To screen which brain regions are obviously affected by HFD in DIO mice, we analyzed c-Fos expression levels in DIO mice. The *in situ* hybrization results indicate PBN had a significantly lower percentage of c-Fos-positive neurons in DIO mice (Fig. 2, A-H). To determine the key molecular markers in the PBN involved in the regulation of sleep and feeding, we treated WT mice with HFD or sleep deprivation and performed RNA-seq analysis in the PBN. The results indicated that *Prokr2* is one of top genes that are upregulated by HFD treatment and sleep deprivation, which indicates *Prokr2* is involved in sleep and bodyweight regulation (Fig. 2, I-K). Next, to better understand the connectivity of PBN with other brain regions, we utilized an HSV-based retrograde tracing method. Our results showed the NPY2R neurons in the LPO extensively projects to the PBN (Fig. 2, L and M). The immunostaining results indicate that ∼50% of LPO neurons projecting to the PBN are NPY2R+ (Fig. 2, N-P and fig. S1).

**Fig. 2.**
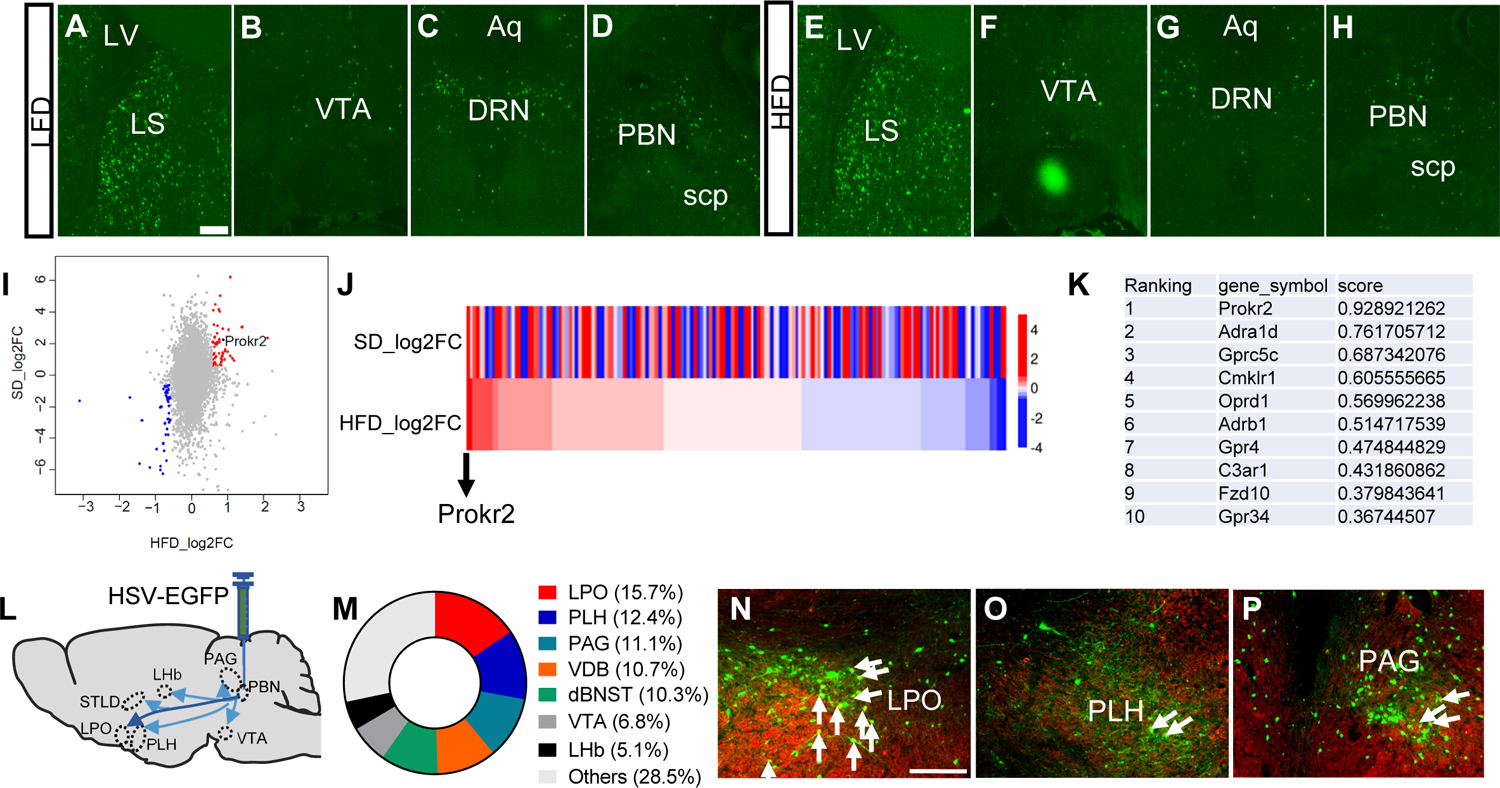
*Prokr2* gene in the PBN neurons is upregulated under the treatment of HFD or sleep deprivation. (**A**-**H**) Representative fluorescence *in situ* hybridization (FISH) image of Fos in the LS (**A**), VTA (**B**), DRN (**C**), and PBN (**D**) in mice with LFD (chow diet) or HFD (**E**-**H**) treatment for 8 weeks. Scale bar in **A** for **A**-**H**, 200 µm. (**I**) Scatter plot analysis of differentially expressed genes of PBN neurons using RNA-Seq analysis under the treatment of HFD or SD. Red and blue dots represent up-or downregulated transcripts, respectively, and gray dots represent transcripts without significant changes under treatment of HFD or SD. (**J**) Heatmap of all the differentially expressed genes from the PBN neurons among *Gpcr* targets genes under the treatment of HFD or SD. (**K**) The top 10 genes upregulated by HFD or SD treatment from **I**. (**L**) Diagram shows retrograde-targeting PBN unilateral injection of *HSV-hEF1*α*-EGFP* into the PBN of WT mice. (**M**) Retrograde targets of PBN in the whole brain. (**N**-**P**) The colocolization of NPY2R and EGFP in the LPO (**N**), PLH (**O**), and PAG (**P**). The NPY2R was stained by immunostaining. Scale bar in **N** for **N**-**P**, 200 µm.

### Activation of PROKR2^PBN^ neurons promoted wakefulness and suppressed food intake

Based on previous studies the PROKR2^PBN^ neurons have high potential to co-regulate sleep and bodyweight (*21, 23, 43–47*). We first investigated the importance of PROKR2^PBN^ neurons in the control of sleep behaviors. Acute optogenetic activation of PROKR2^PBN^ neurons robustly promoted the transition from NREM to wakefulness (Fig. 3, A-C). The duration of NREM and REM sleep episodes also significantly decreased during photostimulation (Fig. 3, D and E). Furthermore, chemogenetic activation of PROKR2^PBN^ neurons also resulted in a decrease of NREM sleep and an increase of wakefulness with a mild effect on REM sleep within 12 hours after CNO injection (Fig. 3, F-J and fig. S2). To test the role of PROKR2^PBN^ neurons in the feeding, we tested the food intake under the optogenetic activation of PROKR2^PBN^ neurons. The results showed a dramatic decrease of HFD food intake during acute photostimulation (Fig. 3K).

**Fig. 3.**
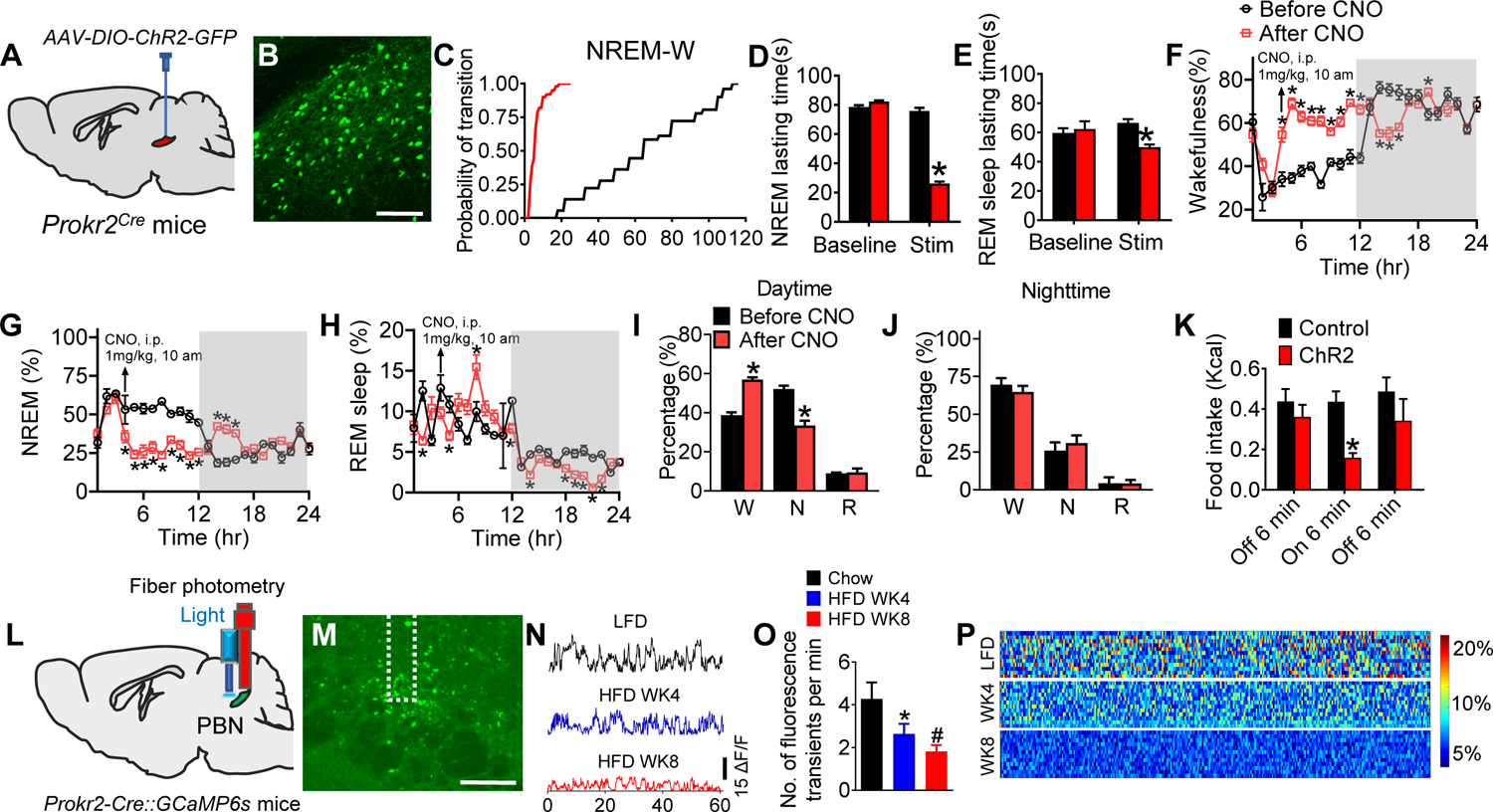
PROKR2 neurons in the PBN are responsible for the comorbidity of obesity and sleep disorder in DIO mice. (**A**) Schematic illustration showing optogenetic activation of PROKR2^PBN^ neurons after injection of *AAV-DIO-ChR2-GFP* into the PBN of *Prokr2^Cre^* mice. (**B**) Representative imaging showing the viral expression of ChR2-EYFP in the PBN. (**C**) Cumulative probability distribution of NREM-to-wakefulness transitions after photostimulation during the daytime period. (**D**, **E**) The duration of NREM (**D**) and REM sleep (**E**) per each episode after photostimulation during the daytime period. (n = 6 per group; *p < 0.05). (**F**-**H**) Time spent in wakefulness (**F**), NREM sleep (**G**), and REM sleep (**H**) during the 24-hr period before and after CNO injection in the *Prokr2^Cre^* mice with *AAV-DIO-hM3Dq-mCherry* into the PBN. The mice were fed with HFD for 8 weeks and the tests were taken on day 56. (n = 8 per group; *p < 0.05). (**I, J**) Total time spent per day in wakefulness, non-REM sleep, and REM sleep in daytime (**I**) and nighttime (**J**) in the mice described in **F**-**H**. (n = 8 per group; *p < 0.05). (**K**) Food intake of HFD before, during and after photostimulation (20Hz/20ms, 6 min) in the mice described in **A**. (n = 9 per group; *p < 0.05). (**L**) Fiber photometry assay revealing the activity of PROKR2^PBN^ neurons of *Prokr2^Cre^* mice with injection of *AAV-FLEX-GCaMP6s* into the PBN. (**M**) *GCaMP6s* was expressed within PROKR2^PBN^ neurons. Scale bar, 200 µm. (**N**-**P**) Representative traces of calcium signals (**N**), fluorescence transients (**O**), and heating map (**P**) of PROKR2^PBN^ neurons in the mice treated with LFD, HFD for 4 weeks and HFD for 8 weeks. (n = 8 per group; *p < 0.05 between chow and HFD WK4, ^#^p < 0.05 between chow and HFD WK8). Error bars represent mean ± SEM. unpaired two-tailed t test in **D**, **E**, **I**, **K**; one-way ANOVA and followed by Tukey comparisons test in **O**; two-way ANOVA and followed by Bonferroni comparisons test in **F**-**H**.

To determine the neural activities of PROKR2^PBN^ neurons during the development of obesity, we analyzed the calcium activities of PROKR2^PBN^ neurons by fiber optometry *in vivo* under HFD treatment. The results showed that PROKR2^PBN^ neurons were progressively inhibited after chronic treatment with HFD (Fig. 3, L-P). Collectively, these results indicate the PROKR2^PBN^ neurons are important in modulating sleep and feeding.

### NPY2R^LPO^ → PROKR2^PBN^ neural circuit regulated sleep and feeding phenotypes

We identified the neural connectivity from NPY2R^LPO^ → PBN in the Fig. 2, to better understand the connectivity of NPY2R^LPO^ neurons with PBN, we utilized ZsGreen-tagged wheat germ agglutinin (WGA), a Cre-dependent trans-synaptic tracer, allowing specific transduction into NPY2R^LPO^ neurons (Fig. 4, A and B). The immunostaining results indicated the NPY2R^LPO^ neurons are innervated with PROKR2^PBN^ neurons (Fig. 4, C-E).

**Fig. 4.**
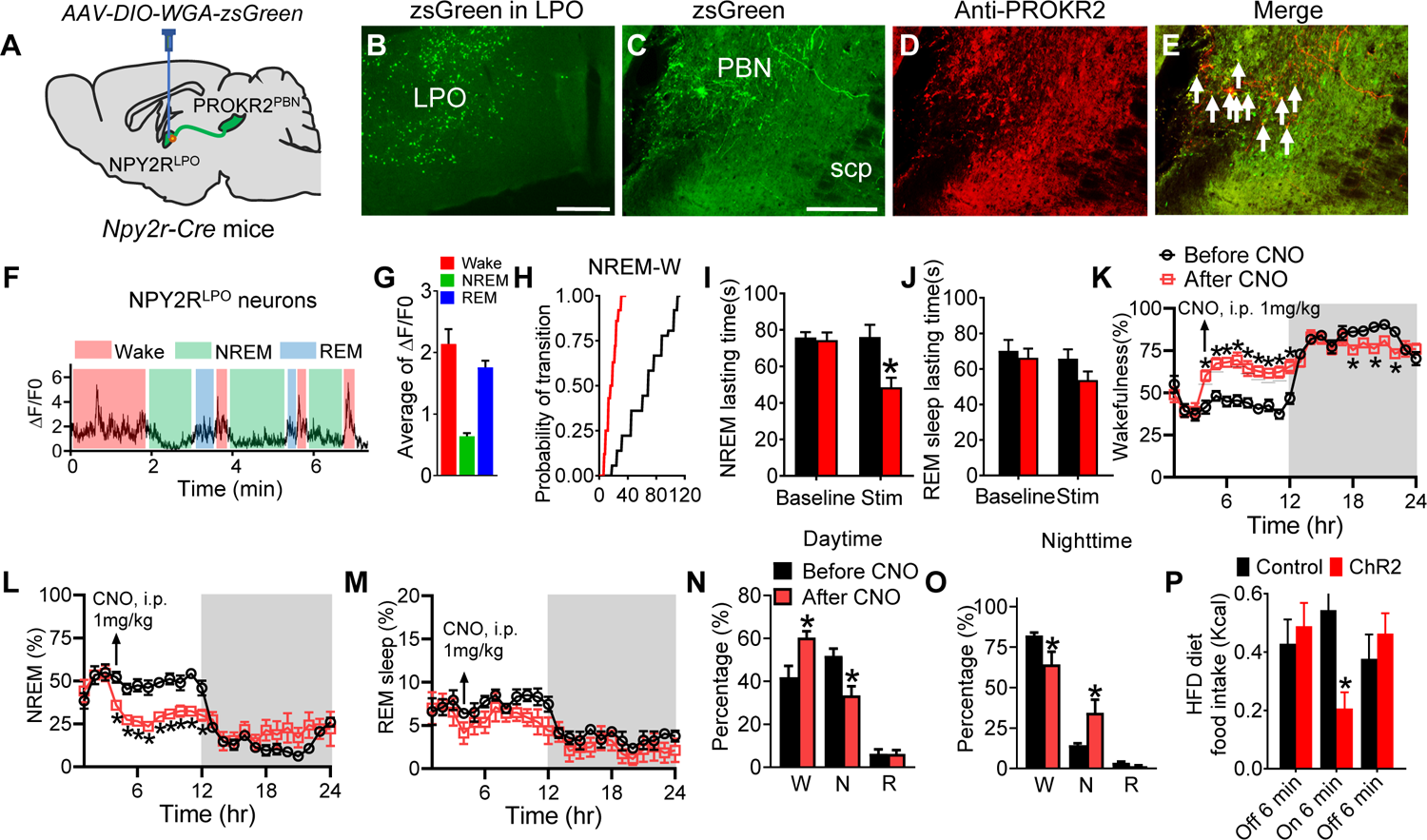
NPY2R neurons in the LPO regulate sleep and feeding behaviors through PROKR2^PBN^ neurons. (**A**) Schematic illustration showing transsynaptic *AAV9-DIO-WGA-ZsGreen* injected into the LPO of Npy2^Cre^ mice. (**B**) Representative images showing WGA-ZsGreen in the LPO. Scale bar, 100 μm. (**C-E**) WGA-ZsGreen-labeled PROKR2 neurons in the PBN with immunostaining of PROKR2. Scale bar in **C** for **C-E**, 100 µm. (**F**) The exemplary calcium imaging of two NPY2R neurons from LPO in a *Npy2r^cre^* mouse with injection of *AAV-FLEX-GCaMP6s* into the LPO. Red shadings indicate brain state as wakefulness, green shadings indicate brain state as NREM, blue shading indicate brain state as REM sleep. (**G**) The average of ΔF/F0 under wakefulness, NREM and REM sleep from NPY2R neurons. (n = 5 per group). (**H**) Cumulative probability distribution of NREM-to-wakefulness transitions after photostimulation during the daytime period by photostimulation of the LPO in the *Npy2^Cre^* mice with the injection of *AAV-DIO-ChR2-EYFP* into the LPO. (**I**, **J**) The duration of NREM (**I**) and REM sleep (**J**) per episode after photostimulation during the daytime period. (n = 6 per group; *p < 0.05). (**K-M**) Time spent in wakefulness (**K**), NREM sleep (**L**), and REM sleep (**M**) during the 24-hr period before and after CNO injection in the *Npy2^Cre^* mice with injection of *AAV-DIO-hM3Dq-mCherry* into the LPO. (n = 8 per group; *p < 0.05). (**N**, **O**) Total time spent per day in wakefulness, NREM sleep, and REM sleep in daytime (**N**) and nighttime (**O**) in the mice described in **K-M**. (n = 8 per group; *p < 0.05). (**P**) Food intake of HFD before, during and after photostimulation of Npy2 fiber in PBN. (n = 6 per group; *p < 0.05). Error bars represent mean ± SEM. unpaired two-tailed t test in **I, J,** and **N-P**; two-way ANOVA and followed by Bonferroni comparisons test in **K-M**.

To determine how the NPY2R^LPO^ neurons respond to the change of sleep and wakefulness, we performed *in vivo* calcium imaging through a gradient refractive index (GRIN) lens coupled to a miniaturized fluorescence microscope. The wakefulness, NREM and REM sleep were recognized based on EEG and EMG recordings. We found that the NPY2R^LPO^ neurons could be regulated by the brain state. The NPY2R^LPO^ neurons showed more active during wakefulness and REM sleep than NREM sleep (Fig. 4, F and G), indicating the neural activities of NPY2R^LPO^ neurons are negatively correlated with NREM sleep.

Next, we studied whether manipulation of the NPY2R^LPO^ → PBN circuit regulates sleep and feeding behavior. Acute optogenetic activation of NPY2R^LPO^ neurons or NPY2R^LPO^ terminals in the PBN robustly promoted the transition from NREM to wakefulness (Fig. 4H and fig. S3, A and B). The duration of NREM and REM sleep episodes also significantly decreased during photostimulation (Fig. 4, I and J and fig. S3, C and D). Chemogenetic activation of NPY2R^LPO^ neurons caused significant sleep dysregulation by increasing wakefulness and decreasing NREM sleep while mildly affecting REM sleep particularly in daytime compared to nighttime (Fig. 4, K-O and Extended Data Fig. 4), which indicated that the activation of NPY2R^LPO^ → PBN circuit induced insomnia symptom. Optogenetic activation of local NPY2R^LPO^ neurons (fig. S3E) or NPY2R^LPO^ terminals in the PBN (Fig. 4P) promoted a robust HFD intake. These results suggest that the NPY2R^LPO^ → PBN circuit is key to regulating physiological sleep and feeding.

### Genetic ablation of *Prokr2* gene in the PBN rescued obese and sleep disorders in DIO mice

To determine the role of *Prokr2* gene in regulating feeding and sleep, we performed site-specific ablation of *Prokr2* in the PBN on base of Cas9/CRISPR. Ablation of *Prokr2* in the PBN obviously improved sleep dysfunctions in DIO mice by restoring sleep and wakefulness time to a normal level in DIO mice (Fig. 5, A-E). Ablation of *Prokr2* in the PBN also decreased food intake of HFD and body weight in DIO mice model (Fig. 5, F-H). Meanwhile, we also observed the typical enhancement of delta power during the transition from wakefulness to NREM after ablation of PROKR2 with less frequent transitions from NREM to wakefulness stages (Fig. 5, I-L). Collectively, we identified *Prokr2* gene that modulates HFD induced obesity and its co-occurring sleep comorbidity.

**Fig. 5.**
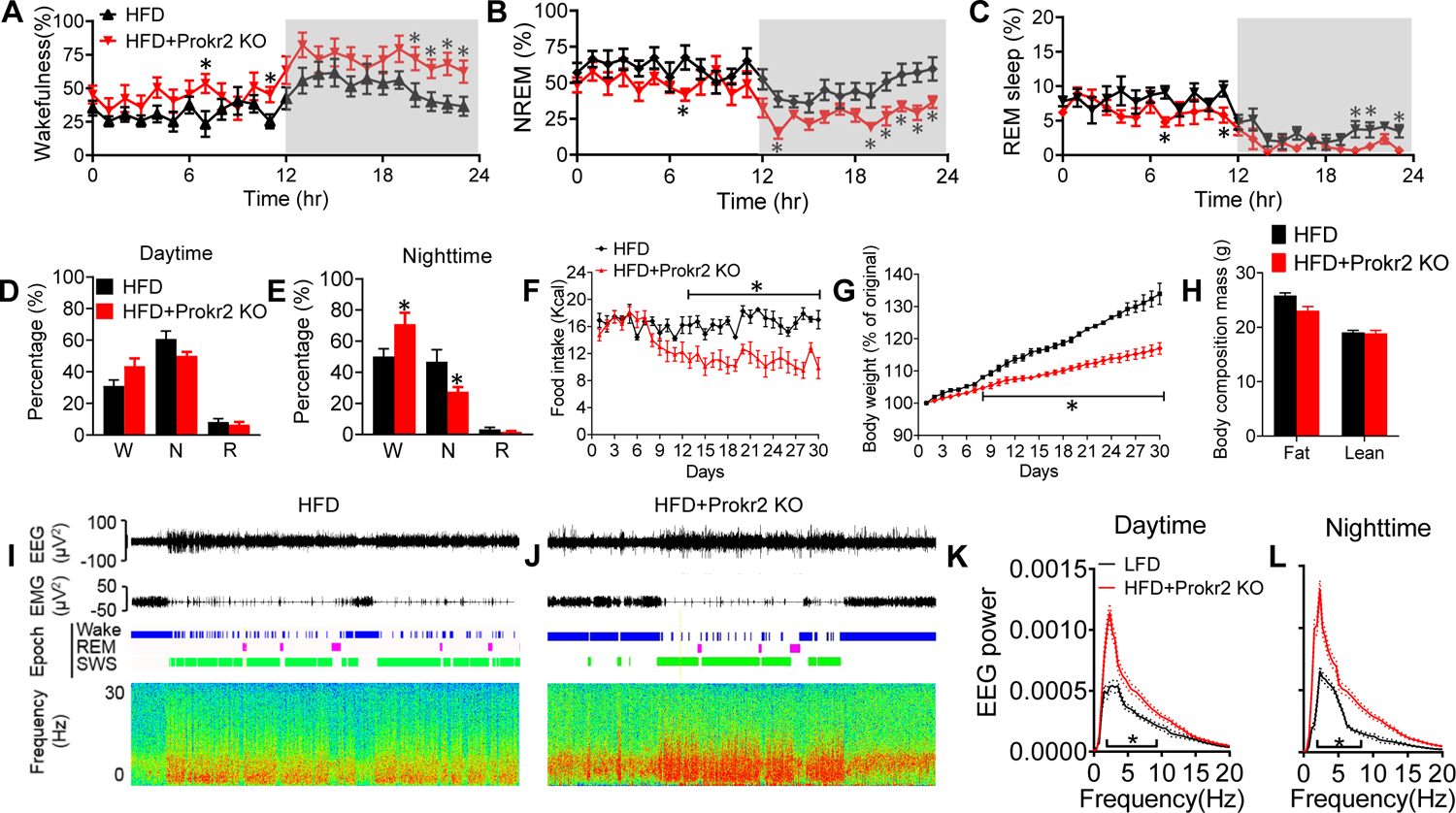
Ablation of PROKR2 improves sleep disorders and obesity in DIO mice model. (**A-C**) Time spent in wakefulness (**A**), NREM sleep (**B**), and REM sleep (**C**) during the 24-hr period on the day 28 in the *Prokr2^Cre^*::*Rosa26^fs-Cas9^* mice injected with *AAV-Prokr2-sgRNA* into the PBN. (n = 6 per group; *p < 0.05). (**D, E**) Total time spent per day in wakefulness, NREM sleep, and REM sleep in daytime (**D**) and nighttime (**E**). (n = 6 per group; *p < 0.05). (**F, G**) Daily food intake (**F**) and body weight (**G**) monitored for 4 weeks in the mice with or without ablation of PROKR2. (n = 6 per group; *p < 0.05). (**H**) Body composition of the mice described in **A, B**. (n = 6 per group; *p < 0.05). (**I, J**) Representative EEG and EMG traces, brain states (color coded), and power spectra from EEG on a 2-hr timescale during daytime in HFD group (**I)** and HFD+PROKR2 KO group (**J**). (**K, L**) EEG power from non-REM sleep during daytime (**K**) and nighttime (**l**) in the mice described in the **A, B**. Error bars represent mean ± SEM. unpaired two-tailed t test in **E**; two-way ANOVA and followed by Bonferroni comparisons test in **A-C, F, G, K, L.**

### Pterostilbene targeted PROKR2 neurons in the PBN to attenuate sleep dysregulations and promote body weight loss

Pterostilbene is a plant-derived stilbenoid, which has been proved to protect the cardiovascular system, relieve obesity and cancer, attenuate diabetes, etc.(*48*). Wondering whether this naturally occurring compound would be a candidate for reversing both feeding and sleep abnormalities, we first tested the effects of pterostilbene on PROKR2 neurons in the PBN in DIO model. *In vivo* fiber optometry revealed that PROKR2^PBN^ neurons are significantly activated after 4th intracerebroventricular (icv) administration of pterostilbene (Fig. 6, A and B). We then tested if pterostilbene could regulate both sleep and bodyweight in DIO mice through icv infusion of pterostilbene. The administration of pterostilbene attenuated the sleep dysfunctions in DIO mice by restoring sleep and wakefulness time to a normal level in DIO mice (Fig. 6, C-I and fig. S5, A-I, and S6). Meanwhile, we also observed the typical enhancement of delta power during the transition from wakefulness to NREM after treatment of pterostilbene with less frequent transitions from NREM to wakefulness stages (Fig. 6, J and K). We further observed pterostilbene induced robust anti-obesity effects on body weight and food intake with a dramatic reduction of fat composition, while lean mass is not altered in all groups, demonstrating there is a significant decrease of fat content, indicating pterostilbene’s fat burning property (Fig. 6, L-N and fig. S5, J and K).

**Fig. 6.**
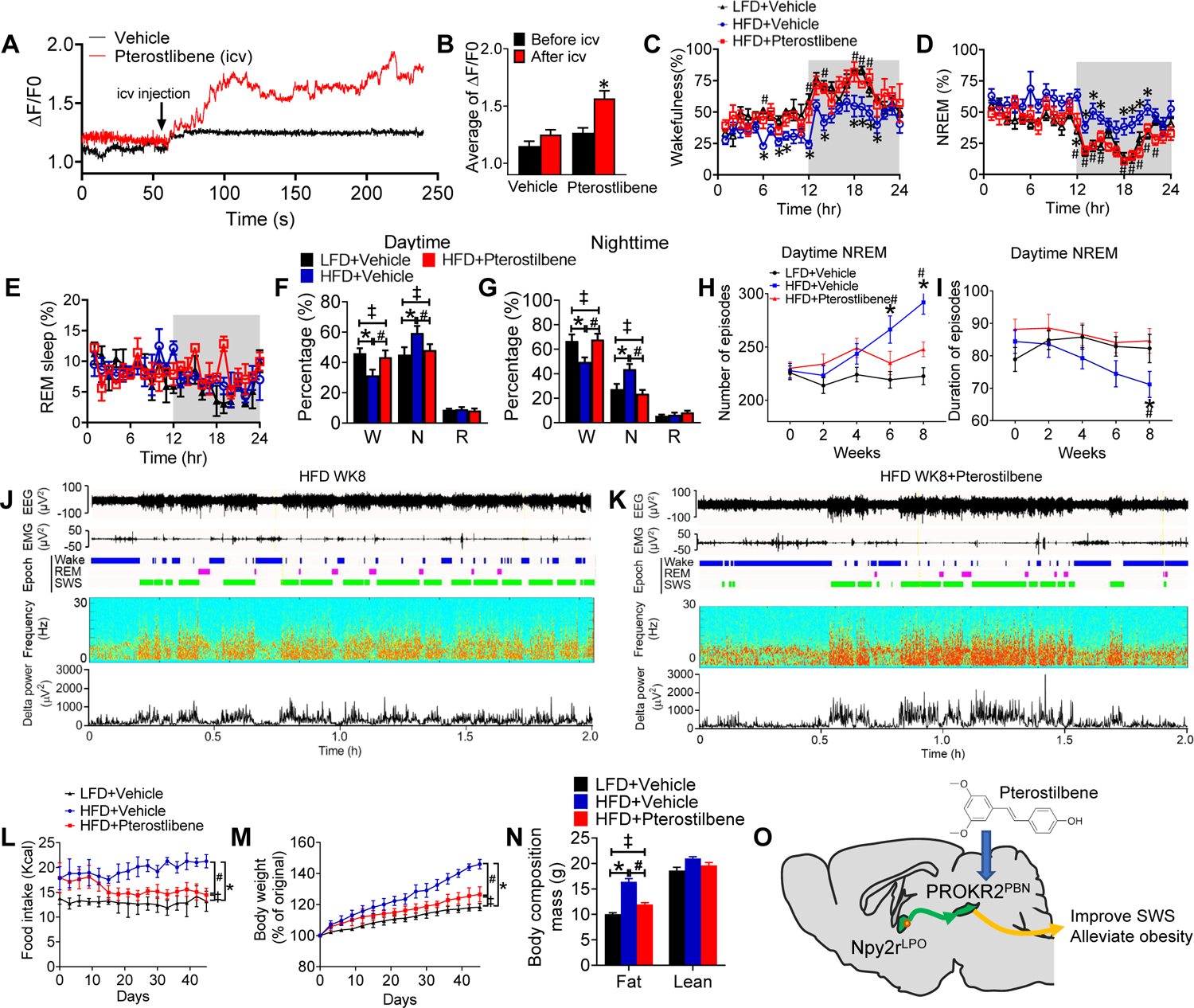
Pterostilbene improves obesity and sleep disorders through targeting PBN. (**A**) The exemplary calcium imaging of PROKR2 neurons from PBN in a *Prokr2^cre^* mouse with injection of *AAV-FLEX-GCaMP6s* into the PBN. (**B**) The average of ΔF/F0 before and after injection of vehicle and pterostilbene by icv from PROKR2 neurons. (n = 5 per group). (**C**-**E**) Time spent in wakefulness (**C**), NREM sleep (**D**) and REM sleep (**E**) during the 24-hr period on the day 56 in the mice fed with LFD, HFD, and HFD+ pterostilbene (10mg/kg, i.p.). (n = 6 per group; *p < 0.05 between LFD and HFD, ^#^p < 0.05 between HFD and HFD+Pterostlibene). (**F**, **G**) Total time spent per day in wakefulness, NREM sleep, and REM sleep in daytime (**F**) and nighttime (**G**). (n = 6 per group; *p < 0.05 between LFD and HFD, ^#^p < 0.05 between HFD and HFD+Pterostlibene, ^‡^p > 0.05 between LFD and HFD+Pterostlibene). (**H**, **I**) The number (**H**) and duration (**I**) of NREM sleep episodes during daytime in the mice described in **C**-**E**. (n = 6 per group; *p < 0.05 between LFD and HFD, ^#^p < 0.05 between HFD and HFD+Pterostlibene). (**J**, **K**) Representative EEG and EMG traces, brain states (color coded), power spectra, and delta power from EEG on a 2-hr timescale during daytime in the mice treated with HFD (**J**) and HFD+ pterostilbene on day 56 (**K**). (**L**, **M**) Daily food intake (**L**) and body weight (**M**) monitored for 8 weeks in the mice described in **C-E**. (**N**) Body composition of the mice described in **C-E**. (n = 6 per group; *p < 0.05 between LFD and HFD, ^#^p < 0.05 between HFD and HFD+Pterostlibene, ^‡^p > 0.05 between LFD and HFD+Pterostlibene). (**O**) A schematic diagram showing a neural circuit in which a subset of NPY2^LPO^ neurons send projections to PROKR2^PBN^ neurons, which reveals the therapy of pterostilbene by action through the PBN, effectively eliminates obesity and sleep disorders comorbidity.

## DISCUSSION

Co-morbidity of obesity and sleep disorder is raising clinical interest, while hardly any evidence has proved or even just mentioned the existence of neural circuit mechanisms. Here, we discovered sleep abnormalities emerged in the early stage of obesity in a DIO mice model. The DIO mice showed progressive sleep fragmentation and excessive low-quality sleep. Through RNA-Seq of PBN neurons from DIO mice and sleep deprived mice, the *Prokr2* gene was selected as top candidate genes that regulate obesity and sleep deprivation simultaneously. With transgenic, optogenetic and chemogenetic tools, we found a neural subpopulation in the PBN, the PROKR2^PBN^ neurons, which could regulate and respond to the intake of HFD. Next, through tracing and immunostaining the dominant upstream target of PROKR2^PBN^ neurons were NPY2R^LPO^ neurons. Furthermore, we deciphered a neural circuit NPY2R^LPO^ → PROKR2^PBN^ that regulated sleep and feeding phenotypes. Ablating PROKR2 signaling in the PBN greatly alleviated diet-induced obesity and hypersomnia. Next, we tested the role of pterostilbene, a stilbenoid drug, in both prevention and treatment of obesity and sleep disorder. It showed pterostilbene ameliorated chronic HFD induced obesity and sleep dysfunctions. Collectively, we established the neural circuit NPY2R^LPO^ → PROKR2^PBN^ is critical for regulating both feeding and sleep (Fig. 6O).

NPY and its receptors are associated with hypothalamic regulation of feeding behavior, metabolism, and energy homeostasis in mammals (*49, 50*). NPY2R-deficient mice exhibited hyperphagia and excessive weight gain (*51*), suggesting an essential role of NPY2R in feeding behavior and body weight control. NPY has also been implicated in the modulation of sleep. NPY plasma levels were increased in patients with obstructive sleep apnea syndrome (*52*). Our results showed that NPY2R^LPO^ neurons are highly correlated with wakefulness and sleep. These neurons are more active during wakefulness and REM sleep, and less active during NREM sleep. So, a central question is how NPY2R neurons modulate numerous complex behaviors and physiological processes including feeding and sleep.

Previous studies identified two GPCRs, PROKR1 and PROKR2, that bind PROK2 with similar affinity in vitro (*53*). PROKR2 was suggested to be the endogenous receptor for PROK2 based on its expression in SCN target regions (*39*). PROK2 overexpression phenotypes could be abolished in PROKR2, but not PROKR1, suggesting that PROKR2 mediates the effects of PROK2 on sleep/wake behaviors. Despite the sleep phenotype observed in PROK2 mutant larvae, it failed to detect sleep defects in either single or double PROKR1 and PROKR2 mutants. This discrepancy suggests there may be additional receptors for PROK2 that are not required for PROK2 overexpression phenotypes but are sufficient to maintain normal sleep/wake behaviors in the absence of PROKR1 and PROKR2. Alternatively, there may be other ligands for these receptors that have effects on sleep opposite to those of PROK2. However, PROKR2 mutants were less active before and more active after lights-on in the morning, consistent with the opposing effects of PROK2 overexpression on behavior in light and dark.

One shining point of our research was to locate a rather small subpopulation of neurons and make it reveal its intriguing functions. PROKR2^PBN^ neurons only took up a minor percentage of all PBN populations and its circuit evidence might very likely be missed or underestimated. Recently, Dan et al. created a genius strategy to find possible sleep regulators brain-wide (*31*). The contradictory part between their and our study was that they didn’t point out the existence of the LPO → PBN inputs, and we think that this was because of the comparably small percentage this subpopulation took up, which would also explain the absence of the LPO → PBN afferent in Luo et al.‘s mapping research (*54*). This situation, we believe, casts us to highlighting the functions of the minority neuron subpopulation and longing for higher-resolution neural tracing techniques.

The PBN’s neural circuit in regulating feeding has been under careful deciphering since 2009, and it’s role in sleep was put up recently by Saper and Fuller et al. (*46*), amending the origin and neural circuit of ascending arousal system (*46, 55*). A small number of works has been released; CGRP^PBN^ neurons promoting arousal during hypercapnia being the most prominent one, which corresponds with Palmiter et al.‘s work (*44*), indicating CGRP^PBN^ as a joint of sleep and metabolism. Our study here points out another subpopulation in the PBN that functions similarly while possessing clinical and epidemiological importance. All these clues remind us that CNS pathophysiological changes may play a pivotal role in various clinical manifestations related to metabolism and sleep, and that the PBN is a very significant joint. There might exist multiple receptors in the PBN sensing metabolic changes across the blood-brain barrier from the periphery and disturbing PBN neural signals, thus resulting in a higher level of appetite, dysregulated metabolic rate, and sleep disorders. Single neuron sequencing to detect whole expression spectrum of every PBN neuron would be a promising way to help tackle this question, although it could be extremely hard to separate adequate single neurons from the tiny PBN and maintain the viability of the vulnerable neurons at the same time.

Pterostilbene has been reported to be effective in animal models of obesity, acting on different metabolic pathways and indicating a pharmacotherapy for the treatment of obesity (*56*). Meanwhile, pterostilbene has preventive effects on the circadian misalignment of mice subjected to sleep restriction. However, the potential mechanism of how pterostilbene improves weight management and sleep remains unknown. An increased understanding of the PROKR2 system can be extended to the treatment of disease beyond obesity because PROKR2 is also implicated in depression, anxiety, and obstructive sleep apnea. These data provide evidence that PBN neurons are a physiologically relevant population of PROKR2–expressing cells involved in the meditation of food intake, body weight reduction and sleep improvement by pterostilbene.

## Acknowledgements

The *AAV-DIO-WGA-ZsGreen* vector was a gift kindly provided by Richard Palmiter. Some AAV vectors were packaged by the Optogenetics and Viral Design/Expression Core at Baylor College of Medicine. Cecilia Ljungberg with the RNA In Situ Hybridization Core facility at Baylor College of Medicine provided technical support on the in situ hybridization.

## Author contributions

Q.W. and Y.H. conceived and designed the experiments. Y.H. and G.X. performed the surgery, EEG/EMG recording, photometry recording, immunohistological imaging, and relevant data analysis. Y.H. performed the isolation of neurons, RNA extraction and RNA-Seq. P.L. and D.G. performed data analysis of RNA-Seq. L.H. performed the colony management, genotyping. Q.W. and Y.H. wrote the manuscript with inputs from all authors on the manuscript. Q.W. and Y.H. supervised the project.

## Data and materials availability

All data needed to evaluate the conclusions in the paper are present in the paper and/or the Supplementary Materials.

---

**Fig. S1. Input of PBN neurons in the brain.** (**A**-**E**) Distribution of retrograde targets of PBN in the whole brain retrograde-targeting PBN by unilateral injection of HSV-hEF1α-EGFP into the PBN of WT mice. The viral expression in the injection site of PBN (**A**), upstream targets in the VDB (**B**), LHb (**C**), dBNST (**D**), and VTA (**E**).

**Fig. S2. The numbers and duration of NREM episodes in daytime and nighttime**. (**A**, **B**) Number of NREM episodes (**A**) and duration of NREM episodes (**B**) during daytime before and after CNO injection in the *Prokr2^Cre^* mice with *AAV-DIO-hM3Dq-mCherry* into the PBN. The mice were fed with HFD for 8 weeks and the tests were taken on day 56. (**C**, **D**) Number of NREM episodes (**C**) and duration of NREM episodes (**D**) during nighttime in the mice described in **A**, **B**. (n = 8 per group; *p < 0.05). Error bars represent mean ± SEM. unpaired two-tailed t test.

**Fig. S3. The effects of optogenetic activation of NPY2^LPO^ neurons on feeding and sleep.** (**A**) Schematic illustration showing optogenetic activation of NPY2^LPO^ neurons after injection of *AAV-DIO-ChR2-EYFP* into the LPO of *Npy2^Cre^* mice. (**B**) Cumulative probability distribution of NREM-to-wakefulness transitions after photostimulation during the daytime period. (**C**, **D**) The duration of NREM (**C**) and REM sleep (**D**) per each episode after photostimulation during the daytime period. (**E**) Food intake of HFD before, during and after photostimulation (20Hz/20ms, 6 min) in the mice described in **A**. (n = 6 per group; *p < 0.05). Error bars represent mean ± SEM. unpaired two-tailed t test.

**Fig. S4. The numbers and duration of NREM episodes in daytime and nighttime.** (**A**, **B**) Number of NREM episodes (**A**) and duration of NREM episodes (**B**) during daytime before and after CNO injection in the *Npy2^Cre^* mice with injection of *AAV-DIO-hM3Dq-mCherry* into the LPO. (**C**, **D**) Number of NREM episodes (**C**) and duration of NREM episodes (**D**) during nighttime in the mice described in the **A**, **B**. (n = 8 per group; *p < 0.05). Error bars represent mean ± SEM. unpaired two-tailed t test.

**Fig. S5. The effects of pterostilbene on sleep and feeding by icv injection.** (**A**-**C**) Time spent in wakefulness (**A**), NREM sleep (**B**), and REM sleep (**C**) during the 24-hr period on the day 56 in mice treated with HFD with or without pterostilbene treatment by icv injection. (**D**, **E**) Percent of time spent per day in wakefulness, NREM sleep, and REM sleep in the mice described in **A** and **B**. (**F-I**) The number (**F**, **H**) and duration (**G**, **I**) of non-REM sleep episodes during daytime and nighttime in mice described in **A**, **B**. (**J**) The change of body weight during the 24-hr period in the mice described in **A**, **B**. (**K**) The food intake of HFD within 24-hr period after icv injection of pterostilbene. (n = 9 per group; *p < 0.05). Error bars represent mean ± SEM. unpaired two-tailed t test in **D-K**; two-way ANOVA and followed by Bonferroni comparisons test in **A-C**.

**Fig. S6. The number and duration of NREM episodes during nighttime in the DIO mice with systematic treatment of pterostilbene.** (**A**, **B**) The number (**A**) and duration (**B**) of NREM sleep episodes during nighttime in mice treated with LFD or HFD with HFD60 with or without pterostilbene. (n = 6 per group; *p < 0.05 between HFD and HFD+Pterostlibene). Error bars represent mean ± SEM. two-way ANOVA and followed by Bonferroni comparisons test in **A**, **B**.

