## Supplementary Figures for "Reversal of Obesity by Enhancing Slow-wave Sleep via a Prokineticin Receptor Neural Circuit"

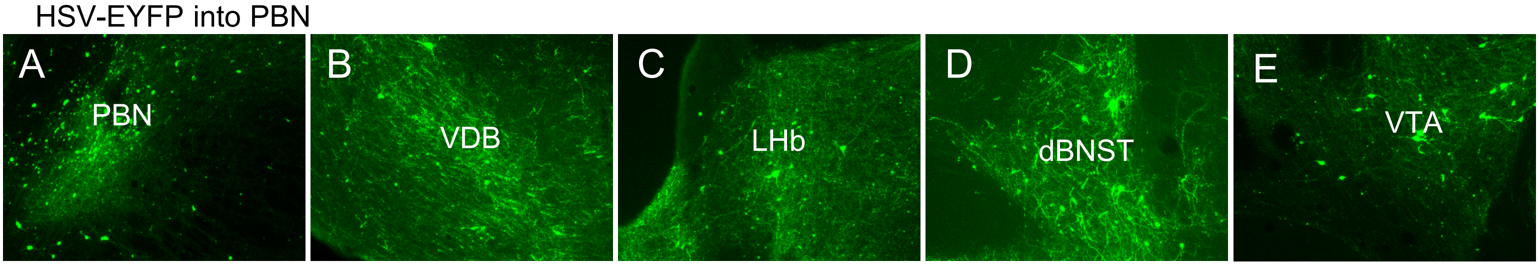


Supplementary Figure 1

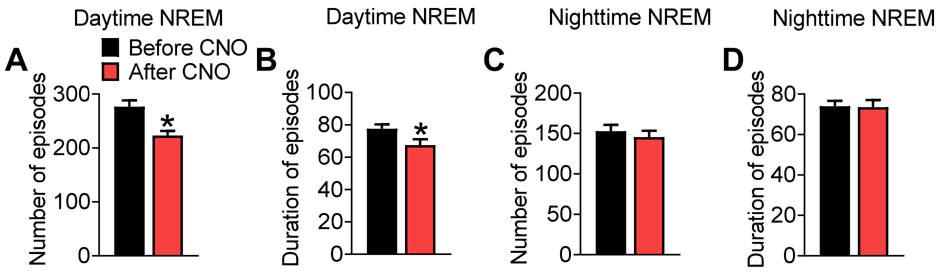


Supplementary Figure 2

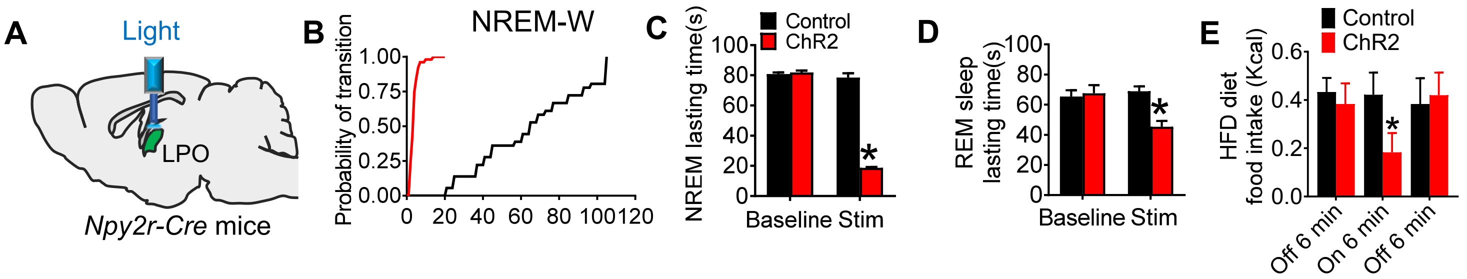


Supplementary Figure 3

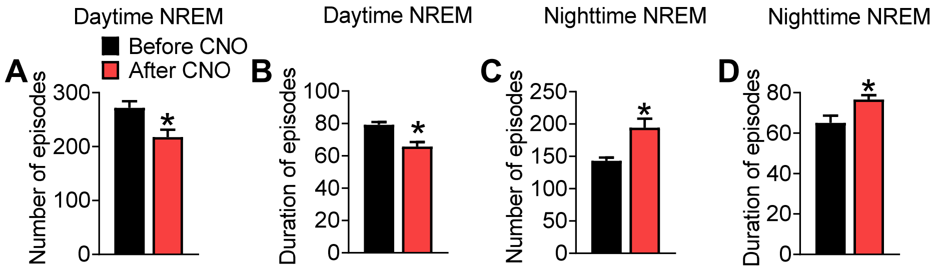


Supplementary Figure 4

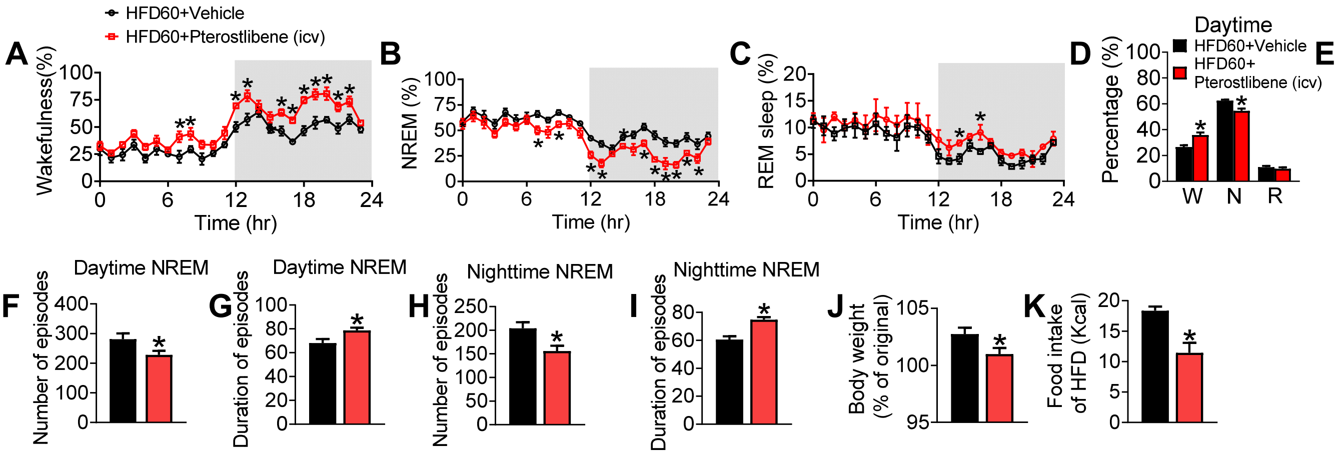


Supplementary Figure 5

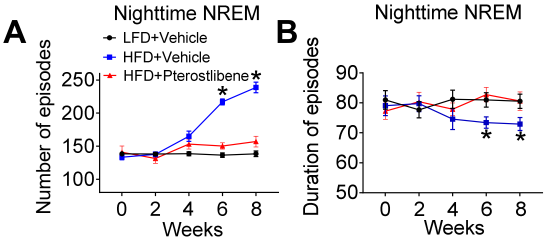


Supplementary Figure 6
